# IBAS: Interaction-bridged association studies discovering novel genes underlying complex traits

**DOI:** 10.1101/2023.08.08.552376

**Authors:** Pathum Kossinna, Senitha Kumarapeli, Qingrun Zhang

**Affiliations:** Department of Biochemistry & Molecular Biology, University of Calgary, Calgary, Alberta, T2N 1N4, Canada; Schulich School of Medicine & Dentistry, University of Western Ontario, London, Ontario, N6A 5C1, Canada; Department of Mathematics and Statistics, University of Calgary, Calgary, Alberta, T2N 1N4, Canada; Alberta Children’s Hospital Research Institute, University of Calgary, Calgary, Alberta, T2N 1N4, Canada

**Keywords:** Genotype-phenotype association study, Stability model, Transcriptome, Gene-gene interactions

## Abstract

The contribution of genetic variants to a complex phenotype may be mediated by various forms of complicated interactions. Currently, the discovery of genetic variants underlying interaction is limited, partly due to that the real interaction patterns are diverse and unknown, whereas exhaustively examining all potential combinations confers the risk of overfitting and instability. We propose IBAS, Interaction-Bridged Association Study, a new model using statistical learning techniques to extract representations of interaction patterns in transcriptome data, which act as a mediator for the next genotype-phenotype association test. Using simulated perturbation experiments, it is demonstrated that IBAS is more robust to noise than similar mediation-based protocols replying on single-genes, i.e., transcriptome-wide association studies (TWAS). By applying IBAS to real genotype-phenotype and expression data, we reported additional genes underlying complex traits as well as their biological annotations. IBAS unlocks the power of integrating gene-gene interactions in association mapping without concerning overfitting and instability.

## Introduction

Discovering genetic basis of complex traits is a long-standing theme in genetics and genomics (Glazier *et al*, 2002). Towards this end, genotype-phenotype association mapping has emerged as an essential tool over the last 20 years (Klein *et al*, 2005; Young *et al*, 2019). In recent times, while identifying genes statistically associated with diseases remains a primary goal, researchers have gradually shifted their focus to their underlying mechanisms through various means of interpretations (Sheng *et al*, 2021). Towards this aim, the availability of multi-scale -omics data especially transcriptome allows researchers to develop sophisticated statistical methods to improve both the discovery of genes and their interpretations (Gamazon *et al*, 2015; Zeng *et al*, 2017; Xie *et al*, 2017; Brandes *et al*, 2020; Okada *et al*, 2016; Xu *et al*, 2017). Among many biologically sensible hypothetical mechanisms, it is intuitive to expect that the variation of gene-gene interaction networks would play a critical role mediating genetic effects to phenotype. Methods such as iPath (Su *et al*, 2021) take this premise and study it at the transcriptomic level, investigating perturbations of given pathways at the sample level and the effect on clinical phenotypes. However, despite gene-gene interactions having been studied for a century (Bateson William *et al*, 1909; Fisher, 1919), no consensus has been reached in understanding the genetic basis of variation of interactions and its implications to phenotypic changes.

As far as our understanding, the lack of statistical genetic models studying the genetic basis of gene-gene interactions is due to the astronomic number of potential combinations, which leads to serious problem of instability. Indeed, studying the interaction patterns associated with phenotypes is already a *p* ≫ *n* problem, suffering from statistical instability as well as computational burden (Fang *et al*, 2019; Zhang *et al*, 2014). By adding an extra layer of the genetic basis of variation of interaction patterns, it only adds substantial risk of instability, potentially leading to many results that may not be robust to perturbation of input data.

Transcriptome-Wide Association Study (TWAS) is a popular model to conduct association mapping utilizing transcriptomes (Gamazon *et al*, 2015; Gusev *et al*, 2016). Although it only focuses on single genes, the in-depth characterization of TWAS by our group (Cao *et al*, 2021a, 2022) offered a unique insight of developing models incorporating interactions. The mainstream format of TWAS is a two-step protocol: first, one can form an expression prediction model using (usually cis) genotypes in a reference dataset (e.g.: GTEx (Consortium *et al*, 2015)). The predicted expression is called Genetically Regulated eXpression, or GReX. Then, in the second step, in the main dataset for association mapping (that does not contain expressions), one can predict expressions using the available genotypes and finally associate the predicted expression to phenotype.

Our recent works showed this interpretation of “predicting expressions” was not the essence of TWAS, but rather, misleading. First, theoretical power analysis showed that TWAS could be more powerful than a hypothetical scenario in which expression data is available in the main GWAS dataset (Cao *et al*, 2021a). Additionally, TWAS could be underpowered compared to GWAS when the expression heritability is low (Cao *et al*, 2021a). Both results question the interpretation of the prediction of expression in TWAS. As such, we proposed to interpret the “prediction” step as a selection of genetic variants directed by expression. From the perspective of Machine Learning, the first step in TWAS is essentially feature selection and the second, feature aggregation. With this interpretation in mind, one may disregard GReX and instead conduct feature selection and aggregation independently. As a result, methodological research focusing on optimal combinations of feature selection and aggregation approaches could be flexibly conducted. Indeed, novel methods splitting these two steps developed by us (Cao *et al*, 2021b, 2022) and independently by others (Tang *et al*, 2021) showed higher power than standard TWAS.

The above unique insight into TWAS has opened a door of conducting association mapping mediated by any “data-bridge” by replacing the gene expression and its predicted value with another entity for feature selection and a form of feature aggregation (Cao *et al*, 2022). Indeed, another development in our group has provided a useful tool leveraging brain images (He *et al*, 2023) in a similar manner.

In this work, we designed an unconventional framework by forming a mediator representing interactions computed using gene expressions through statistical learning techniques. We then used it to facilitate the discovery of the genetic basis of variations of interactions and their consequence to phenotypic changes. More specifically, starting from a gene expression matrix of all genes in a prespecified pathway, we extract representative statistics such as PCA (Hotelling, 1933), t-SNE (der Maaten & Hinton, 2008) or UMAP (McInnes *et al*, 2018) to form the objective function for feature selection. Then, following our previous work (Cao *et al*, 2021b, 2022; He *et al*, 2022), a kernel machine (Wu *et al*, 2010) is employed for feature aggregation aimed at association test. The outcomes are single genes whose genetic variants alter the variation of gene-gene interactions in the focal pathway, which also impact the phenotypic changes. This method is named “Interaction Bridged Association Study”, or IBAS. Importantly, such representative statistics contributed by many genes in the pathway is more robust to noise than single genes alone, therefore intuitively could be more stable mediators than single genes in a typical TWAS protocol. Indeed, perturbation experiments using GTEx data show that IBAS is more stable than standard TWAS protocols that rely on single genes. We further applied IBAS to the seven diseases of the Wellcome Trust Case Control Consortium, or WTCCC (The Wellcome Trust Case Control Consortium, 2007) and discovered novel genes underlying pathways and diseases.

## Results

### The IBAS Framework

Step 1 of IBAS utilizes a reference dataset where gene expression and genotype data are available. Known biological pathways (or even user specified collection of genes known to contain interactions) are then used to extract the expression data. The extracted subset of gene expressions (corresponding to a particular pathway) then undergoes dimensionality reduction (using methods such as PCA, t-SNE or UMAP) to produce interaction components (“data-bridges”) **(Fig 1A**). These components are expected to capture interactions at the transcriptome level. In <u>Step 2</u>, each component is then associated with individual *cis*-variants corresponding to the genes in the pathway (**Fig 1B**). The coefficients obtained for each variant indicate their contribution to the interactions captured by the corresponding interaction component. <u>Step 3</u> of IBAS involves a GWAS dataset where genotype data is assessed together with a phenotype of interest. Variants in the GWAS dataset are then aggregated to the gene level, weighted by their contribution to interactions (obtained in Step 2 depicted in **Fig 1B**) and tested for association with phenotypes using the Sequence Kernel Association Test, or SKAT (Wu *et al*, 2010) (**Fig 1C**). This results in the identification of genes associated with the phenotype of interest through the mediation of the pathway being considered.

**Figure 1.**
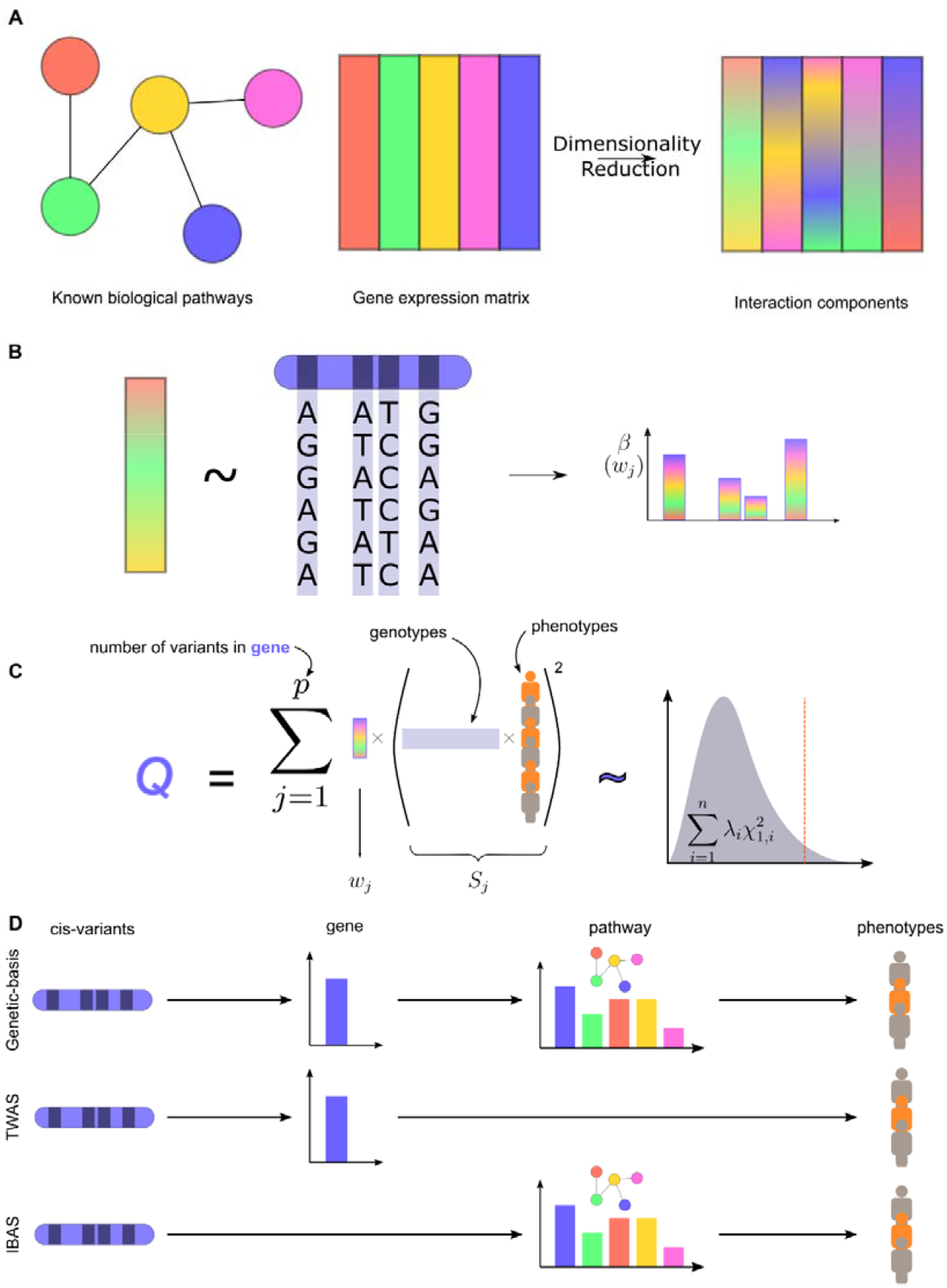
Overview of IBAS Framework. **A)** Gene expression matrices of a reference dataset (such as GTEx) are subset for known biological pathways (from sources such as KEGG, GO, etc.) and undergo dimensionality reduction (using methods such as t-SNE and UMAP) to produce interaction components (“data-bridges”). **B)** Individual interaction components are then associated with *cis*-variants of the genes in the pathway. The coefficients (or significance) of association are interpreted as the contribution of the variants towards the interaction effects at the transcriptome level. **C)** Genotypes in a GWAS dataset are then tested for association with a phenotype of interest, aggregated at the gene level using the Sequence Kernel Association Test, weighted by the interaction contributions obtained in (B). **D)** The natural cascade of genetic effects affecting phenotype are modelled to various extents by existing methods. TWAS methods capture the effect of variants on individual genes’ expression and ignore the contributions of interactions at the pathway level on phenotype while IBAS captures the effect of variants on these interactions (where the individual gene contribution may be also present).

The natural cascade of genetic variants affecting phenotype may be depicted in **Fig 1D, upper panel**: from variants to gene expression, to pathways and finally phenotype. TWAS captures the effects of variants on gene expression, and then gene expression on phenotype, ignoring the contribution of interactions at the pathway level (**Fig 1D, middle panel**). IBAS on the other hand, captures the effects of cis-variants on the interactions at the pathway level (where it could be argued that the gene-level effect itself is present) and identifies association with the phenotype (**Fig 1D, lower panel**). Therefore, IBAS reveals the genetic basis of complex traits from a compensatory angle.

### IBAS is more stable than single gene-mediated TWAS

To test whether interaction-based protocol is more stable than single-gene based ones, perturbation experiments were conducted to compare the stability of IBAS to PrediXcan with respect to different reference data sets (**Fig 2**). We utilized real data from the GTEx Consortium for building the data-bridge and investigate the effects on the Type 1 Diabetes (T1D) cohort from the WTCCC dataset (along with the other 6 cohorts in the **Appendix Fig S1-S6** to ensure this stability holds across different genetic architectures). To simulate various signal-to-noise ratios, expression data in the GTEx dataset was perturbed by adding uniform noise of varying ranges (= 0.1, 0.25 and 0.5) relative to the expression levels of each gene (**Materials & Methods**) to produce 10 replicates each for every noise-level. PrediXcan elastic-net models were trained on these data sets, and IBAS weights were obtained for dimensionality reduced data-bridges of the same. These models, and weights were then used for association analysis in WTCCC T1D data. In the case of IBAS, for each dimensionality reduction method, weights were combined for all components by mean for coefficient-based and Fisher combined p-values for the p-value based method (we call this combination ‘meta’-weights) (**Materials & Methods**).

**Figure 2.**
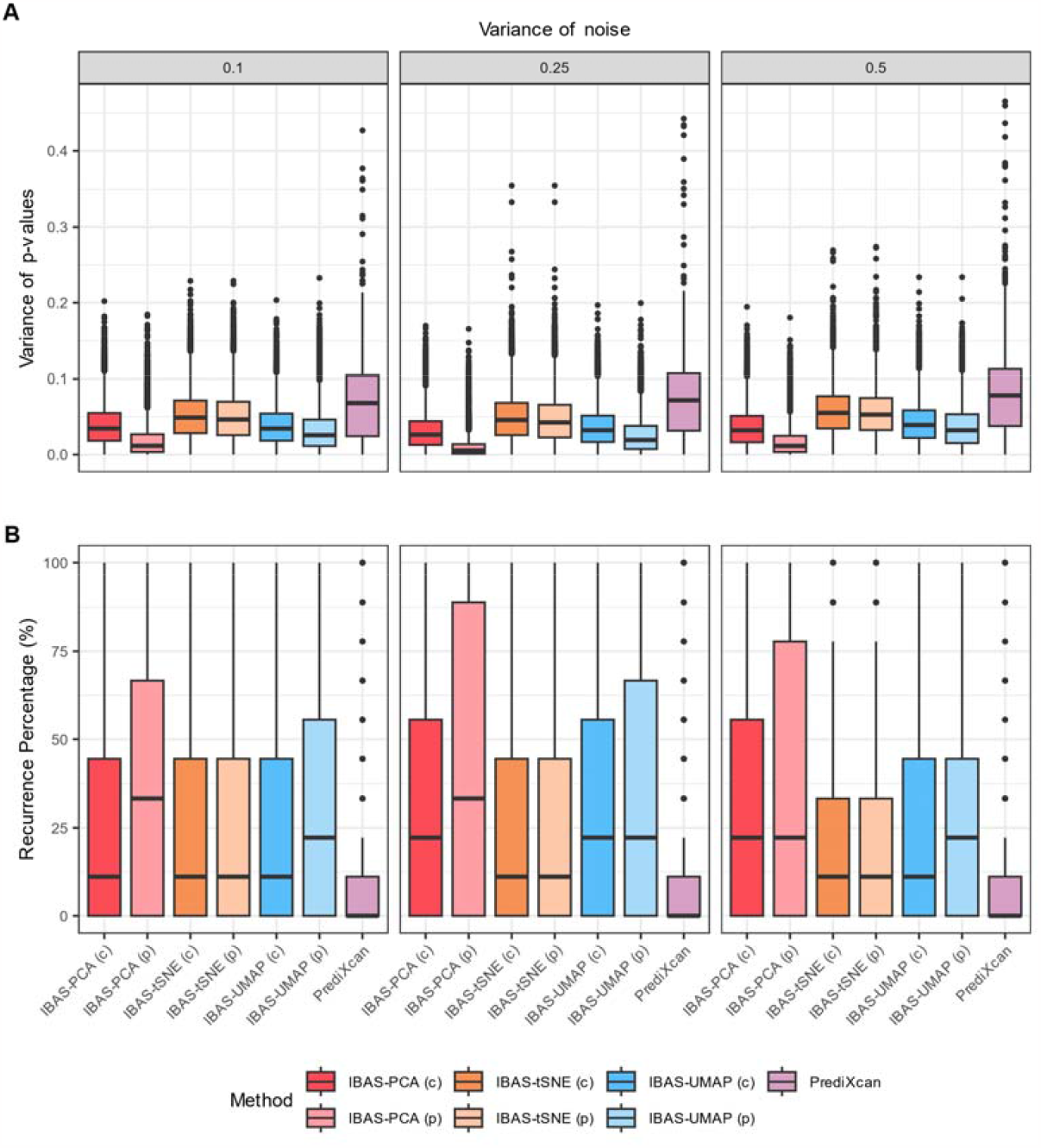
Simulated Perturbations reveal robustness of IBAS to noise in reference data. Simulations were conducted by perturbing gene expression values of each gene by percentage ranges (0.1, 0.25 and 0.5) in the reference data. 10 perturbed expression datasets (replicates) were generated for each of the noise levels and variability of results for case-control association with the Wellcome Trust Case Control Consortium Type 1 Diabetes cohort was evaluated (other cohorts were tested and available in **Appendix Fig S1-S6**) using both IBAS and PrediXcan. PCA, t-SNE and UMAP dimensionality reduction methods as well as p-value based (p) and coefficient based (c) interaction association weights were tested in the case of IBAS. **A)** Indicates the variance of observed p-values in genes identified as significant () in at least one replicate while **B)** indicates the recurrent significance of genes identified as significant in at least one replicate.

IBAS-based methods showed clear advantages over PrediXcan in stability of discoveries. **Fig 2A** showcases that IBAS has lower variance in p-values over a wide range of noise levels for genes identified as significant (*p* < 0.05) in at least one replicate. PCA-based IBAS demonstrates a significantly lower variability in p-values. It is also notable that weighting by p-value, rather than by coefficient across all dimensionality reduction methods reduces variability of p-values. We also look at the recurrence percentage (**Fig 2B**) of genes that were significant in at least one replicate and uncover a similar pattern. PrediXcan performs poorly indicating that a gene uncovered in one replicate may not be uncovered again if there was even a slight perturbation in the reference data. IBAS is significantly more stable – with PCA-based, p-value weighted IBAS performing the best (**Fig 2A**).

### IBAS discovers novel genes through the mediation of interactions

IBAS uncovered significant discoveries across the 7 diseases of the WTCCC case-control cohort; a number of which were previously identified in the DisGeNET database (Piñero *et al*, 2017) with an additional number of discoveries that were novel (**Fig 3**). Most discoveries were for the Rheumatoid Arthritis (RA) and T1D cohorts both of which also contain the most annotations in the DisGeNET database being autoimmune diseases that have been extensively studied. However, IBAS was also able to uncover significantly associated genes for Bipolar Disorder (BD), Coronary Artery Disease (CAD), Hypertension (HT) and Type-2 Diabetes (T2D). Furthermore, the discovery of numerous novel genes is expected and welcomed since most association studies incorporate purely genetic information (or, in the case of TWAS, transcriptomic information of a single gene).

**Figure 3.**
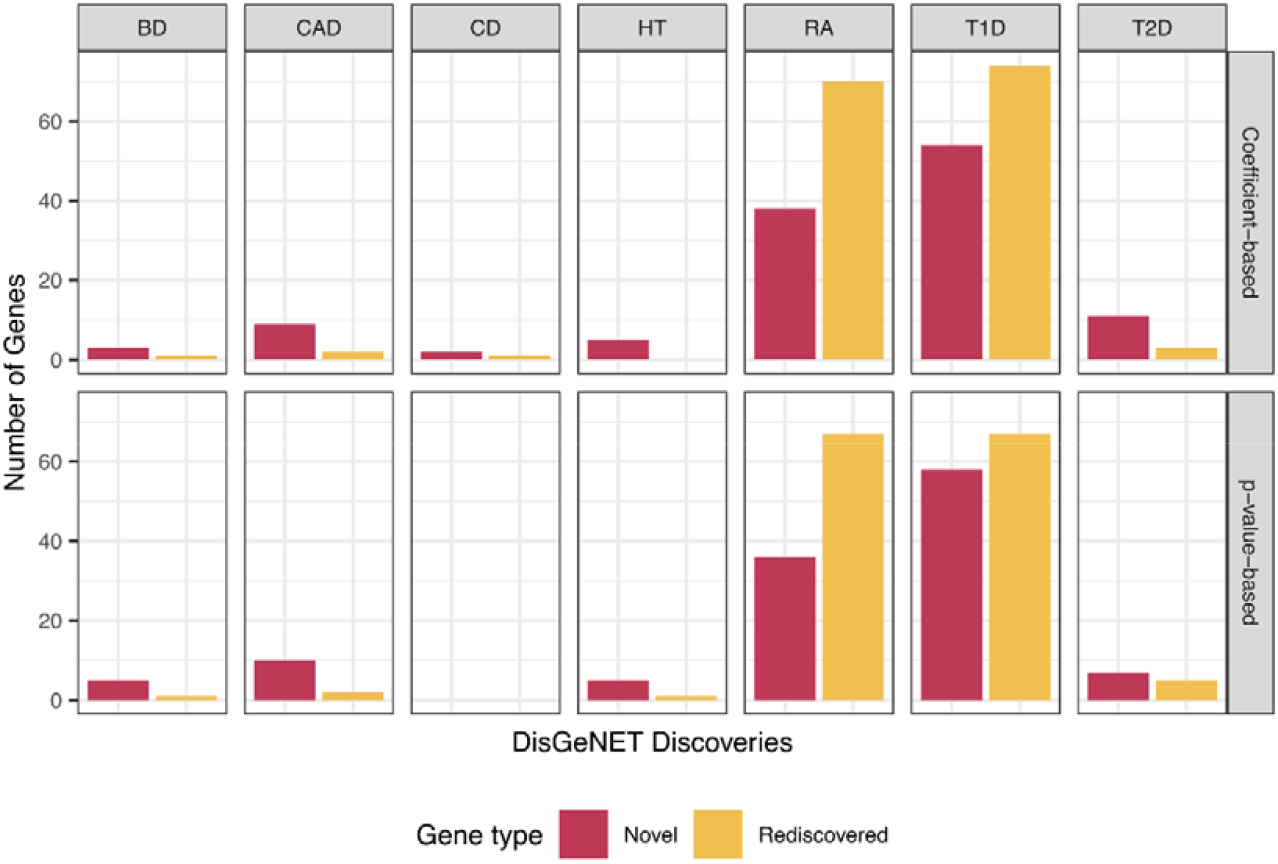
Evaluation of genes associated with disease uncovered by IBAS. Genes identified as associated with each of the 7 diseases (columns) in the WTCCC dataset (with GTEx used as reference) across both coefficient-based and p-value based (rows) for the PCA data-bridge were annotated as “Rediscovered” if found as previously associated with the same disease in the DisGeNET database. Remaining genes are annotated as “Novel”. Evaluation of t-SNE and UMAP can be found in **Appendix Fig S7-S8**.

IBAS shows contrastive results compared to other gene-disease association tools (**Fig 4**). PrediXcan (Gamazon *et al*, 2015) and kTWAS (Cao *et al*, 2021b) are used for this comparison since they represent analyses based on the typical TWAS framework and a kernel-based framework similar to IBAS but without the interaction effects, respectively. Results from IBAS are once again based on the meta-weights (mean for coefficient based and Fisher combined p-values for p-value based across all dimensionalities reduced components). kTWAS and PrediXcan were found to report fewer results in most diseases, but with a level of overlap between them that suggests that while kTWAS does gain more power due to the use of the kernel-based test for feature aggregation as shown previously (Cao *et al*, 2021b), they extract similar information from transcriptomic data. IBAS on the other hand, incorporates interactions from gene expression and thus is expected to discover genes based on their association with these interactions. The high number of genes discovered in T1D and RA by both kTWAS and PrediXcan further support the theory previously mentioned that there may be a high marginal effect from genes associated with these diseases. It is also noteworthy that IBAS uncovers several genes associated with HT: one of which, *CLOCK*, has known associations with HT in the DisGeNET database (**Fig 3**). This further strengthens the hypothesis that IBAS is able to effectively uncover genes that may be associated with disease through their interactions at the pathway level.

**Figure 4.**
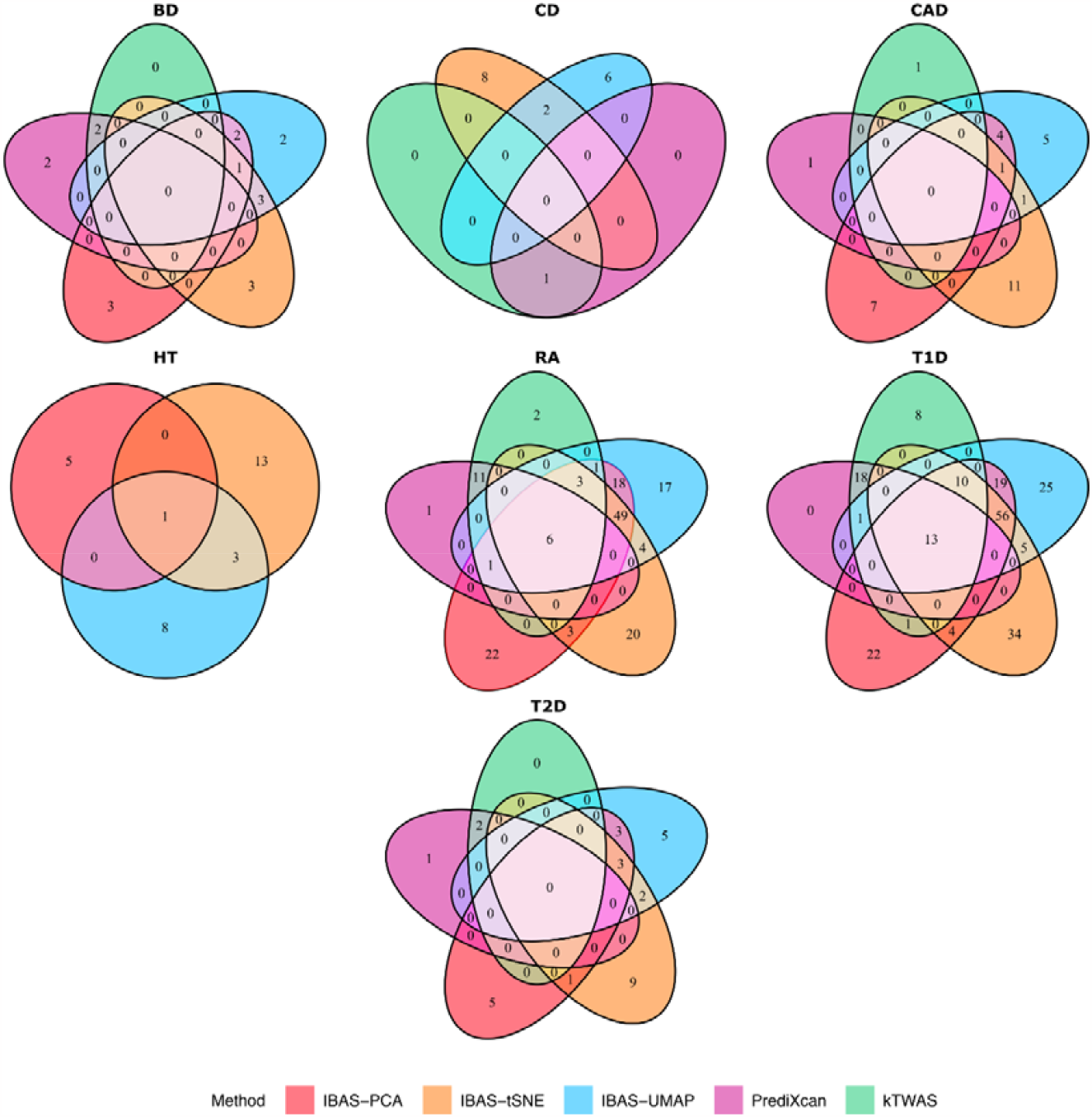
Comparison of IBAS results with kTWAS and PrediXcan. Genes identified by each of the IBAS methods (PCA, t-SNE and UMAP) as well as PrediXcan and kTWAS were compared to identify overlap and uniqueness of discoveries across the 7 diseases of the WTCCC dataset.

### Biological annotation of results

IBAS distinguishes itself from previous annotations in the WTCCC cohort by identifying genes that may have a small marginal effect but contribute to disease phenotype mediated through relevant pathways (**Fig 5**). In particular, observing the patterns in diseases where many significant genes are identified such as RA and T1D, previously annotated diseases are identified across multiple pathways – indicating that their strong marginal effect may have contributed to their significance. This could explain the discovery of a large number of genes in the HLA region being identified as significant for these auto-immune disorders (Serrano *et al*, 2006) as well as *TNF* (Serrano *et al*, 2006) and *ITPR3* (Huang *et al*, 2010). The weaker marginal effects of genes may explain the lower number of genes identified to be associated with the other diseases since most methods do not account for pathway-level interactions as IBAS does.

**Figure 5.**
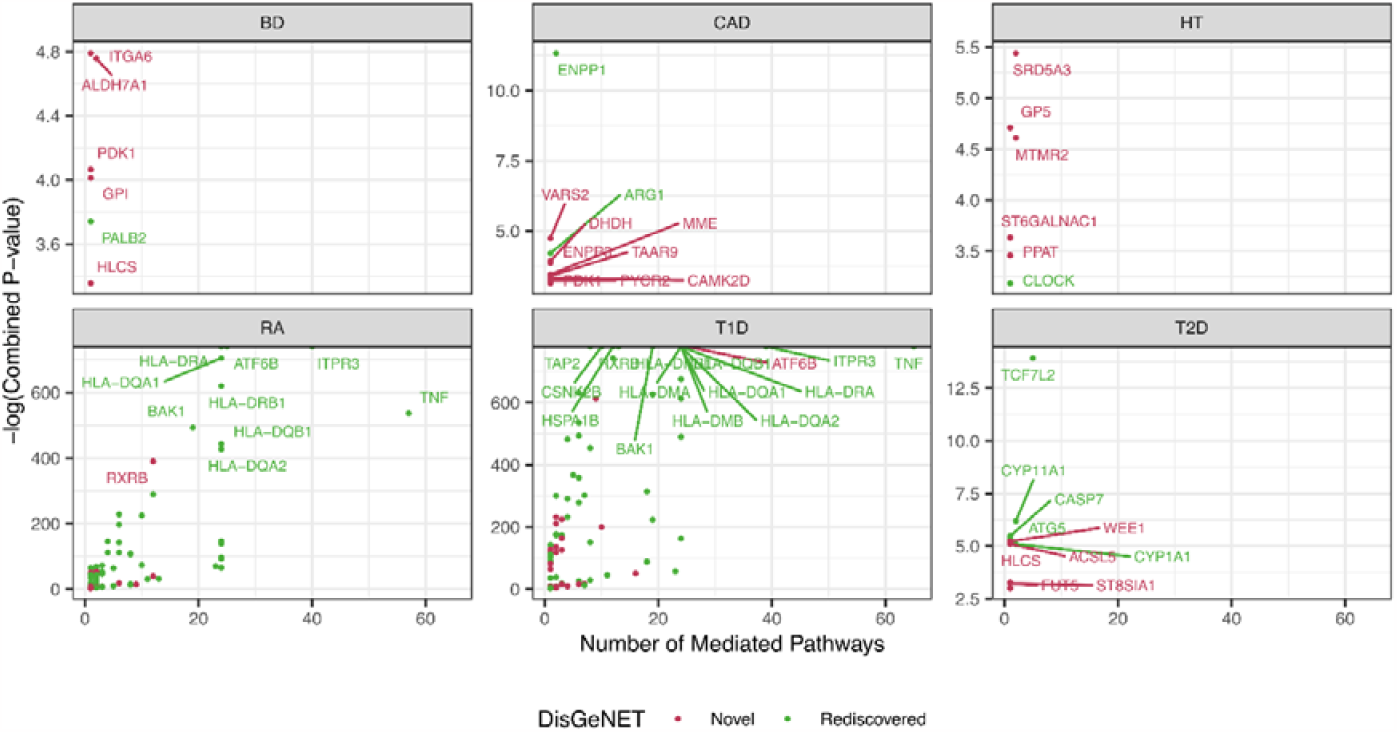
IBAS identifies associations with marginal effects as well as interaction-driven effects. Comparison of significance of previously discovered (“Rediscovered”) and “Novel” genes associated with the 6 disease (no significant genes associated with CD using IBAS-PCA) cohorts of the WTCCC dataset. Combined P-values (y-axis) indicate Fisher combined p-values of genes identified across multiple pathways; while the x-axis denotes the number of pathways mediated through which the gene was identified in as associated with the phenotype.

IBAS identifies a key mechanism suggesting the mediation of metabolism related pathways and genes in BD. 4 out of the 5 novel genes identified by IBAS are associated with various metabolic processes (GPI Gene - GeneCards | G6PI Protein | G6PI Antibody; Yang *et al*, 2019; Dupuy *et al*, 2015; Zempleni *et al*, 2014). The role of metabolism in psychiatric disorders has long been investigated (Zuccoli *et al*, 2017) with a clear association established between dysfunctional metabolism and psychiatric disorders including BD (Rosso *et al*, 2015).

*HLCS*, a gene known for its role in biotin metabolism (Zempleni & Kuroishi, 2012) in the body, is found to be highly associated with BD through the biotin metabolism pathway (Kanehisa Laboratories; Kanehisa & Goto, 2000) by IBAS indicating a potential pathway that could mediate the development of BD. Biotin or Vitamin B7 is known to be vital to brain function as it is a crucial component in the process of glucose metabolism (Kennedy, 2016). Furthermore, *HLCS* is highly specific to neurons (both inhibitory and excitatory) in the brain according to the Human Protein Atlas (Karlsson *et al*, 2021; The Human Protein Atlas). Changes in the excitatory and inhibitory (E/I) synaptic balance have been previously associated with bipolar disorder in animal models (Lee *et al*, 2018) and the specificity of HLCS as well as its importance in metabolism make it a potential mediator in the emergence of the BD phenotype. While no studies have been conducted on the direct association of *HLCS* on BD, one study identified increased methylation at CpG sites in *HLCS* in borderline personality disorder patients in their genome-wide association study but failed to find the same in a validation analysis (Teschler *et al*, 2013). Increased methylation could indeed be a potential mechanism of repression of *HLCS* expression, resulting in dysfunctional biotin-dependant metabolic pathways which in turn could create an imbalance in the activation of E/I neurons contributing to manic episodes.

In the case of other significant associations, IBAS hints at potential biologically pivotal candidates in disease etiology. *VARS2*, the most significant gene identified with association to CAD (excluding all previously annotated genes in DisGeNET) has been identified to cause heart failure in zebrafish models when knocked-out (Kayvanpour *et al*, 2022). Another significant gene, *ENPP3*, has known cardiac-specific function and is downstream of *NKX2-5*, a transcription factor associated with congenital heart disease (Barth *et al*, 2010). *PPAT*, identified to be significant in the HT phenotype, was concluded to be an important gene in the control of BP rhythm in rats (Murata *et al*, 2020). A significant gene (*ACSL5*) identified in T2D, has been studied to be the primary target of the most strongly associated variant of T2D in the *TCF7L2* gene (Xia *et al*, 2016), and is important in the metabolism of fatty acids.

## Discussion

### Stability and validations

We expect that the development of IBAS is a significant progress towards the goal of developing statistically stable models involving interactions. This assumption is because IBAS does not aim to explore all combinations of interactions and therefore does not run the risk of overfitting at this stage. Indeed, we have revealed that the performance of IBAS is stable with respect to the alternation of input expression data (**Fig 2**). Additionally, as t-SNE and UMAP can extract nonlinear components incorporating interactions flexibly, its performance should be also stable to alternations of the prior knowledge of membership of pathways.

More rigorous validations are needed to quantitatively support the above assumptions. First, numerical simulations with known genetic architectures must be conducted to assess the theoretical properties of IBAS. This is not a trivial task as there is very limited literature reporting patterns of gene-gene interactions that involve many genes in a pathway. During the development of IBAS, we have carried out simulations following the three non-linear genetic architectures, i.e., epistatic, compensatory, and heterogenous models. These non-linear models are better accepted by the field and have been used in the previous publications in our group (Cao *et al*, 2021b, 2022; Li *et al*, 2020). However, these architectures are designed to mimic interactions of two genes and are therefore not suitable for the purpose of verifying IBAS. Instead, we opted to use perturbations of real data in our simulations since there was a lack of literature and benchmarking surrounding the simulation of realistic pathway-level interactions.

### Association v.s. Causality

It is worth noting that, although IBAS used PCA/t-SNE/UMAP represented gene-gene interactions to mediate the association test, we are not claiming the causality of “genetics ⇒ interaction patterns ⇒ phenotype”. Instead, there could be all kinds of causality models, for instance, the pleiotropic model in which genetics causes changes in both interaction patterns and phenotype. Here, the key insight is that we used the interaction patterns to methodologically mediate the discovery, despite the real biological causality relationship is unknown. The same issue applies to many existing works leveraging multi-scale omics to identify association between genetics and phenotype (Gamazon *et al*, 2015; Zeng *et al*, 2017; Xie *et al*, 2017, 2016; Cao *et al*, 2021b, 2022; Brandes *et al*, 2020; Okada *et al*, 2016; Xu *et al*, 2017).

## Materials and Methods

### IBAS Implementation (I): Biological pathways

Biological pathways refer to known interactions among genes, proteins and metabolites (van Iersel *et al*, 2008). These interactions are often curated from literature by various projects such as Kyoto Encyclopedia of Genes and Genomes (KEGG) (Kanehisa *et al*, 2017, 2023; Kanehisa & Goto, 2000), Reactome (Joshi-Tope *et al*, 2005) or WikiPathways (Kelder *et al*, 2012). These databases provide access to both downloading and visualizing these pathways and their known interactions. This work utilizes the KEGG database through the use of the KEGGREST (Tenenbaum & Maintainer, 2022) R package. However, IBAS could be applied in a similar manner to any of the pathway databases or even other databases containing known biological interactions such as the Gene Ontology Resource (Consortium, 2019).

### IBAS Implementation (II): Dimensionality Reduction

The downloaded pathways provide context for sub-setting the reference data, but we must turn to dimensionality reduction to uncover the complex interactions within the pathways. Here, we take advantage of three methods often used in the analysis of genomics data: PCA (Hotelling, 1933), t-distributed Stochastic Neighbour Embedding (t-SNE) (der Maaten & Hinton, 2008) and Uniform Manifold Approximation and Projection (UMAP) (McInnes *et al*, 2018). These methods are based on the concept of identifying a low-dimensional representation of points in a high-dimensional space. We use these methods to reduce the dimensionality of the pathways to a maximum of 10 components in the case of PCA and UMAP, and 2 in the case of t-SNE. In instances where a pathway has less than 10 genes, we set the number of components to be generated as the number of genes.

***PCA*** utilizes the principle of iteratively identifying the combinations of components of the higher-dimensional space that maximize captured variance. These components are identified in such a way that they are uncorrelated with all previous PCs resulting in orthogonal combinations of the original data (Lever *et al*, 2017). These components have been identified in various fields to capture variance and identify differences in subpopulations of the data (Reich *et al*, 2008; Jollife & Cadima, 2016).

***t-SNE*** was originally designed for the purpose of visualizing high-dimensional data in a low-dimensional space (typically 2-dimensional). t-SNE is now often used in the visualization of transcriptomic data, particularly in RNA-Seq (Kobak & Berens, 2019). t-SNE is based upon the concept of minimizing the probability distribution of points in a low-dimensional space close to their probability distribution in a high-dimensional space. Consider a set of points ***x***_*i*_ where ***i*** ∈{**1**,…,***n***} and *x*_*i*_ ∈ **ℜ**^***p***^ and their embedding in a lower-dimensional space ***y***_*i*_ where ***y***_*i*_ ∈ **ℜ^*d*^** and ***d*** « ***p***.

The conditional probability between two distinct points ***x***_*i*_ and ***x***_*i*_ in the higher-dimensional space is defined as:

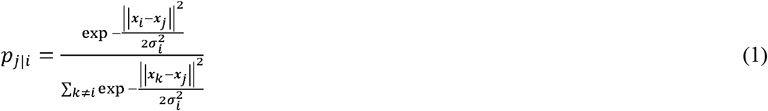

and *p*_*i*|*i*_= 0. We then define the joint probability of the two points ***x***_*i*_ and ***x***_*j*_ as:

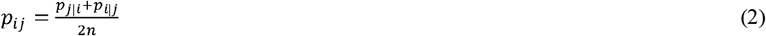

The joint probability of the same points in the lower dimension is then modelled using a Student t-distribution with one degree of freedom as:

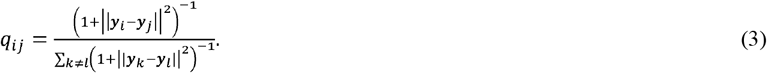

The Kullback-Leibler (KL) divergence between P and Q distributions is then used as the cost function and optimized via gradient descent to obtain the optimal embedding.

***UMAP*** uses concepts similar to t-SNE, to achieve a low-dimensional embedding. The mathematics behind UMAP are fairly complex (McInnes *et al*, 2018), and for the purposes of this paper are left out and instead, a high level explanation provided. UMAP begins by constructing what is termed a “fuzzy simplicial complex” which is a type of weighted graph between the points in the high-dimensional space with the edges denoting the likelihood that two points are connected. To determine this connectivity (similar to **Equation 1**), UMAP extends a radius determined by the distance to the nth nearest neighbour. Then, the likelihood is reduced based on the distance from the point. It is also required that each point is connected to at least its closest neighbour. The last step involves constructing a similar network to the first, but this time in the lower dimensional space, with the use of cross-entropy as the cost function compared with the KL-divergence used by t-SNE.

### IBAS Implementation (III): Interaction Directed Feature Selection

Once a pathway has undergone dimensionality reduction, it is then in a form suitable for association analysis to “select” relevant genetic variants (here we reference feature selection with regards to feature prioritization). Using the same reference dataset, cis-variants are tested for association with each individual component of the interaction components. Each component is analyzed individually as it is expected that different components would capture a different set of interactions among the gene expression in the pathway. While any number of methods could be used for this association analysis, we use the linear mixed model (Kang *et al*, 2008, 2010) which accounts for population stratification by the use of a kinship matrix (which is also called a Genomic Relationship Matrix (GRM) in some literature).

The linear mixed model is defined as:

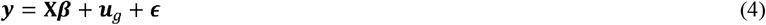

where ***y*** denotes the vector of phenotypes (interaction component in IBAS), **X** is the matrix containing the intercept, variant to be tested and other covariates,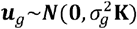 denotes the random effects and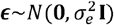. Here **K** denotes the kinship matrix which provides a measure of the degree of relatedness of the individuals in the sample.

This association is carried out for all cis-variants of the genes in the pathway. While it is possible that trans-variants also do influence the expression of the genes in the pathway, such testing is extremely resource-intensive and would suffer from a large multiple-testing problem (Goeman & Solari, 2014). Typically, cis-variants are considered as the variants within 1Mb of the start and end of the gene and we used this definition for our analysis as well.

### Genotype-Phenotype Association Analysis via Feature Aggregation

Finally, gene-level testing is conducted using the Sequence Kernel Association Test (SKAT). This test combines the effect of multiple variants in a kernel-based score test to test phenotypic association of the aggregate:

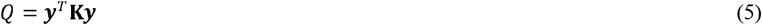

where y is the vector of phenotypes and **K** is a kernel function based on the centralized genotype matrix of the GWAS dataset, **G**. While multiple kernels may be defined for the use in the score test (Wu *et al*, 2010), we use the modified kernel previously used in our work (Cao *et al*, 2021b, 2022):

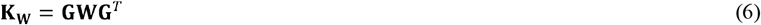

where**W**=diag(*α*_1_,…,*α*_*i*_,…, *α*_*p*_). *α*_*i*_ here, denotes the coefficients of association derived from **Equation 4** for the reference data. These coefficients of association may be selected as the direct coefficients from the linear mixed model (i.e. *α*_*i*_ = |*β*_*i*_|; coefficient-based weighting) or as the p-values associated with each coefficient from the same (i.e. *α*_*i*_= - log *p*_*i*_; p-value based weighting). This is then used in **Equation 5** to calculate Q which follows a mixture of chi-square distributions (Davies, 1980) (**Fig 1C**).

While weights associated with individual components of each dimensionality reduction method could be used to identify genes effecting phenotypes through various interactions, a ‘meta’-weight was also considered for each pathway. In this case, the weights for each variant were calculated as the mean of the coefficients in the case of coefficient-based weighting, and in the case of p-value based weighting, the Fisher combined p-values. These ‘meta’-weights could be assumed as the average contribution of each variant on the major interactions in the pathway under study.

Thus, the outputs of IBAS are genes associated with phenotypes of interest, mediated by the interaction effects of the pathway considered. Since a single pathway may contain up to hundreds of genes, a multiple hypothesis testing correction in the form of a Benjamini-Hochberg correction (Benjamini & Hochberg, 1995) is applied to reduce the risk of false positives.

### Other Methods compared to IBAS

#### TWAS: PrediXcan

PrediXcan (Gamazon *et al*, 2015) trains an elastic-net (Zou & Hastie, 2005) predictive model of gene expression levels on reference data in a similar manner to IBAS as described before. Considering the expression of gene *j* to be denoted as *T*_*j*_, the PrediXcan model can be stated:

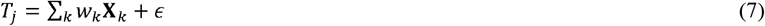

where *X*_*k*_ denote the (*cis-*)variants of gene *j*. Once the model is trained, it is used to predict the Genetically Regulated eXpression (GReX) in a GWAS dataset. Finally, the predicted expressions are associated with the phenotypes of interest using either linear, logistic or Cox regression models or other non-parametric methods (such as testing for Spearman’s correlation) (Gamazon *et al*, 2015).

#### kernel-based TWAS (kTWAS)

kernel-based TWAS (kTWAS) (Cao *et al*, 2021b) was the first data bridge model proposed by our group. Following the first steps of PrediXcan, an elastic-net model is trained on reference data. However, instead of using this model to predict gene expression in the GWAS dataset, the weights from the elastic-net model, **W**_*kTWAS*_ = diag(*γ*_1_,…, *γ*_*i*_,… *γ*_*p*_), are used in SKAT in the form of a weighted kernel matrix for the score test. Similar to IBAS, these weights may be derived from the elastic-net model used in PrediXcan as *γ*_*i*_ = *w*_*i*_ or using weights derived from p-values associated with each term. Following the results in the original paper, we use the coefficient-based analysis for comparison. Essentially, the kernel matrix **K**_w_ is constructed as:

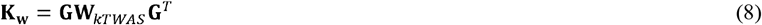

and the score statistic *Q* is defined as:

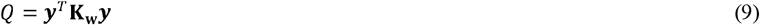

which is distributed as a mixture of chi-squared distributions (**Fig 1C**) (Davies, 1980).

### Real Data Analysis: Reference Data

Publicly available, normalized, filtered expression data of “Whole Blood” tissue from the GTEx Consortium (Consortium *et al*, 2015) which contains samples from 670 individuals and for 20315 transcripts was used as the reference expression data. This data was then corrected for PEER factors (Stegle *et al*, 2010), the top 5 genotype principal components, age and gender as suggested by the GTEx Portal. This expression data was then used for dimensionality reduction by subsetting for individual pathways.

Genotype data for 838 individuals and 43,066,422 SNPs were present in the raw VCF data. Since IBAS works best if there is a high degree of overlap between the GWAS and reference datasets when considering genotypes, imputation was carried out using SHAPEIT4 (Delaneau *et al*, 2019) for phasing and IMPUTE5 (Rubinacci *et al*, 2020) for the final imputation based on data from the 1000 Genome Project (Consortium & others, 2015). This results in 88,863,451 SNPs for the imputed data. The data was then filtered to include only individuals who had expression data available and general QC (MAF <0.05, deviation from HWE *p*<10^-8^ and relatedness > 0.025) was applied, resulting in 6,187,683 SNPs for 553 individuals. This data was converted to the {tped, tfam} file format and JAWAMix5 was used to convert it to hdf5 (Long *et al*, 2013) for out-of-core association analysis with the interaction components obtained from the expression data.

### Real Data Analysis: GWAS Data Containing Genotype and Phenotype

The Wellcome Trust Case Control Consortium (The Wellcome Trust Case Control Consortium, 2007) contains genotype data for 13,241 cases and 2,938 controls (individual sample sizes are provided in **Appendix Table S1**) across 7 diseases: bipolar disease (BD), coronary artery disease (CAD), Crohn’s disease (CD), rheumatoid arthritis (RA), type 1 diabetes (T1D), type 2 diabetes (T2D) and hypertension (HT). The original dataset only contained 393,272 variants since this data was obtained using the Affymetrix GeneChip 500K arrays and thus imputation (Rubinacci *et al*, 2020) was conducted as in GTEx genotype data to produce 73,162,888 variants.

### Simulations Generating Perturbated Data

Simulations were carried out by adding perturbations to gene expression data from the Whole Blood tissue in GTEx (Consortium *et al*, 2015) to evaluate robustness to changes in reference data. Each expression value was regenerated using a uniform distribution *U*~(*x*(1+*r*), *x*(1-*r*)) where *x* is the current expression value and *r* = {0.1, 0.25, 0.5}. This approach attempts to hold the same expression patterns within pathways while adding individual noise levels to genes as could be present due to various factors such as sampling time and sequencing batch effects. The varying levels of *r*, adds increasing levels of noise and 10 replicates are generated at each level. These 30 individual expression datasets are then used in conjunction with unperturbed genotype data in generating data-bridges for IBAS and in training PrediXcan models for association testing with WTCCC cohorts.

## Author contributions

Designed the study: PK, QZ

Implemented the tool: PK

Analyzed data: PK, SK, QZ

Wrote the manuscript: PK and QZ

## Data and materials availability

GTEx gene expression: https://gtexportal.org/home/datasets

GTEx whole genome sequencing data: https://www.ncbi.nlm.nih.gov/projects/gap/cgi-bin/study.cgi?study_id=phs000424.v9.p2

WTCCC genotype: https://www.wtccc.org.uk/

IBAS source code available at: https://github.com/QingrunZhangLab/IBAS

