## Appendix (Supplementary Materials) for "IBAS: Interaction-bridged association studies discovering novel genes underlying complex traits"

Appendix Figure S7 – Evaluation of genes associated with disease uncovered by IBAS-UMAP. 9

Appendix Figure S8 – Evaluation of genes associated with disease uncovered by IBAS-tSNE. 10

### Appendix Table S1 – Sample sizes of WTCCC cohorts

| **Disease** | **Case** | **Control** |
|---|---|---|
| Bipolar Disorder (BD) | 1868 | 2938 |
| Coronary Artery Disease (CAD) | 1926 | 2938 |
| Crohn's Disease (CD) | 1748 | 2938 |
| Rheumatoid Arthritis (RA) | 1860 | 2938 |
| Type 1 Diabetes (T1D) | 1963 | 2938 |
| Type 2 Diabetes (T2D) | 1924 | 2938 |
| Hypertension (HT) | 1952 | 2938 |

**Appendix Figure S1 – Simulated Perturbations reveal robustness of IBAS to noise in reference data (BD).**

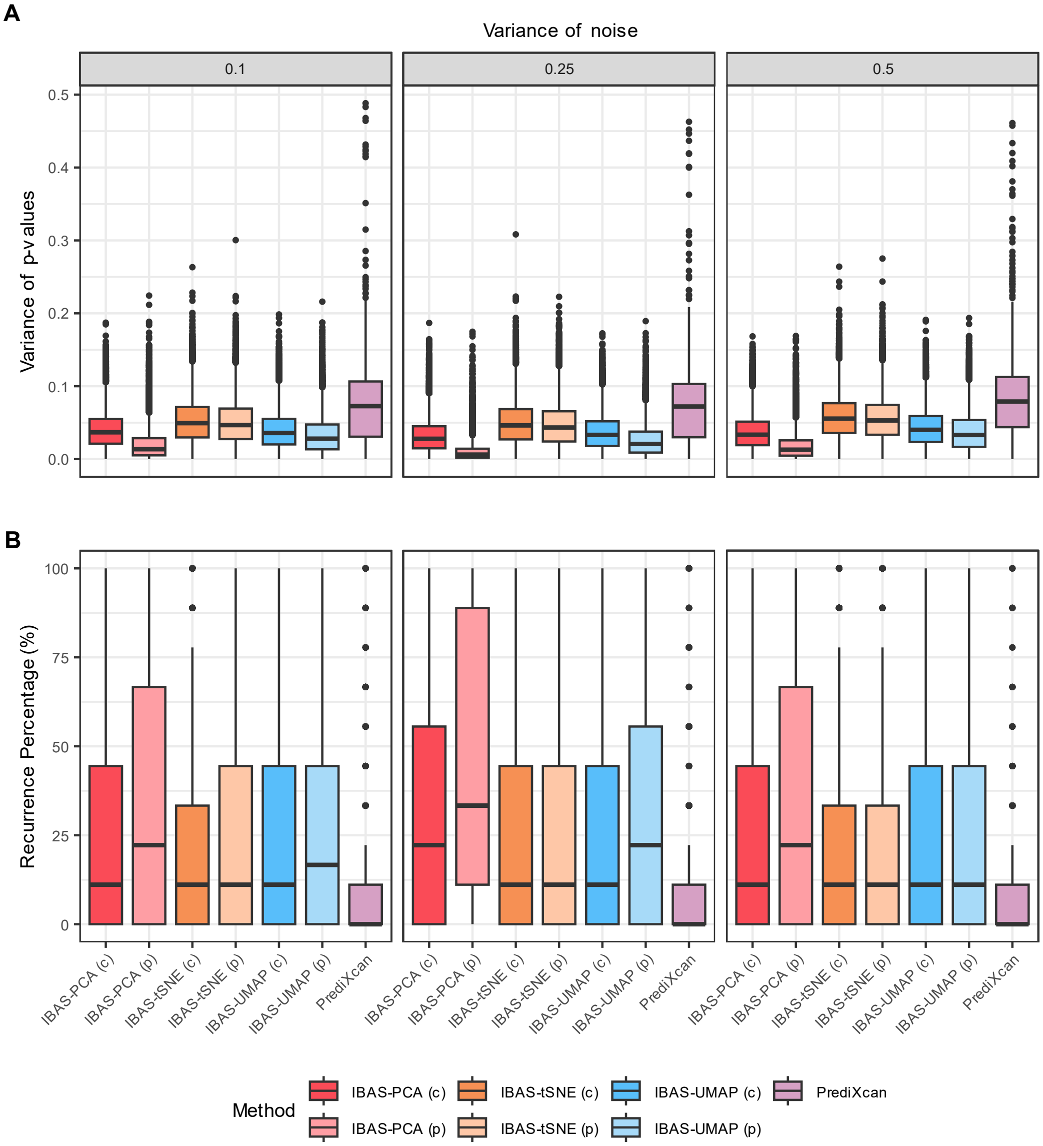

©

Simulations were conducted by perturbing gene expression values of each gene by percentage ranges (0.1, 0.25 and 0.5) in the reference data. 10 perturbed expression datasets (replicates) were generated for each of the noise levels and variability of results for case-control association with the Wellcome Trust Case Control Consortium Bipolar Disorder cohort was evaluated using both IBAS and PrediXcan. PCA, t-SNE and UMAP dimensionality reduction methods as well as p-value based (p) and coefficient based (c) interaction association weights were tested in the case of IBAS. **A)** Indicates the variance of observed p-values in genes identified as significant ($p < 0.05$) in at least one replicate while **B)** indicates the recurrent significance of genes identified as significant in at least one replicate.

**Appendix Figure S2 – Simulated Perturbations reveal robustness of IBAS to noise in reference data (CAD).**

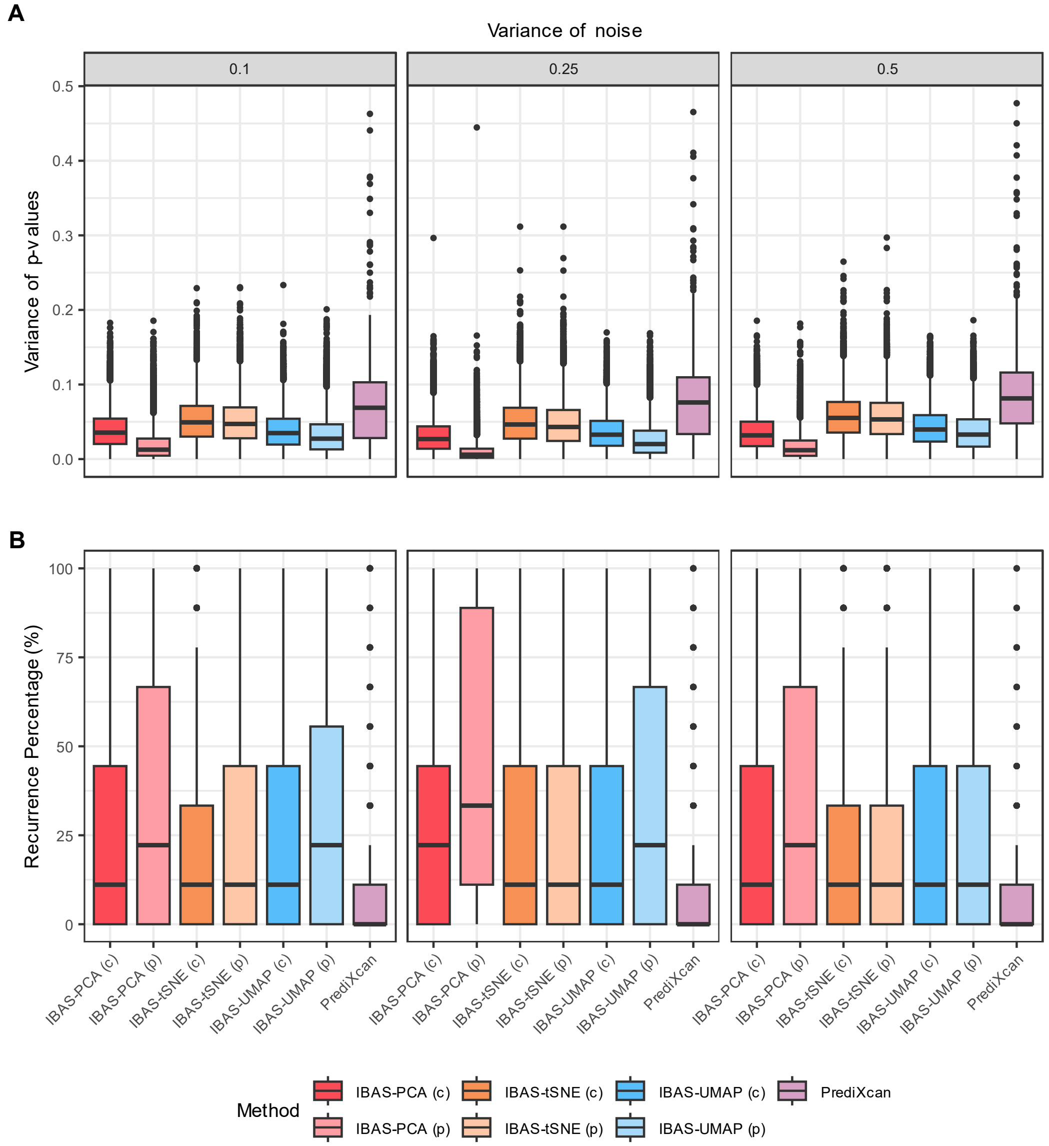

Simulations were conducted by perturbing gene expression values of each gene by percentage ranges (0.1, 0.25 and 0.5) in the reference data. 10 perturbed expression datasets (replicates) were generated for each of the noise levels and variability of results for case-control association with the Wellcome Trust Case Control Consortium Coronary Artery Disease cohort was evaluated using both IBAS and PrediXcan. PCA, t-SNE and UMAP dimensionality reduction methods as well as p-value based (p) and coefficient based (c) interaction association weights were tested in the case of IBAS. **A)** Indicates the variance of observed p-values in genes identified as significant ($p < 0.05$) in at least one replicate while **B)** indicates the recurrent significance of genes identified as significant in at least one replicate.

**Appendix Figure S3 – Simulated Perturbations reveal robustness of IBAS to noise in reference data (CD).**

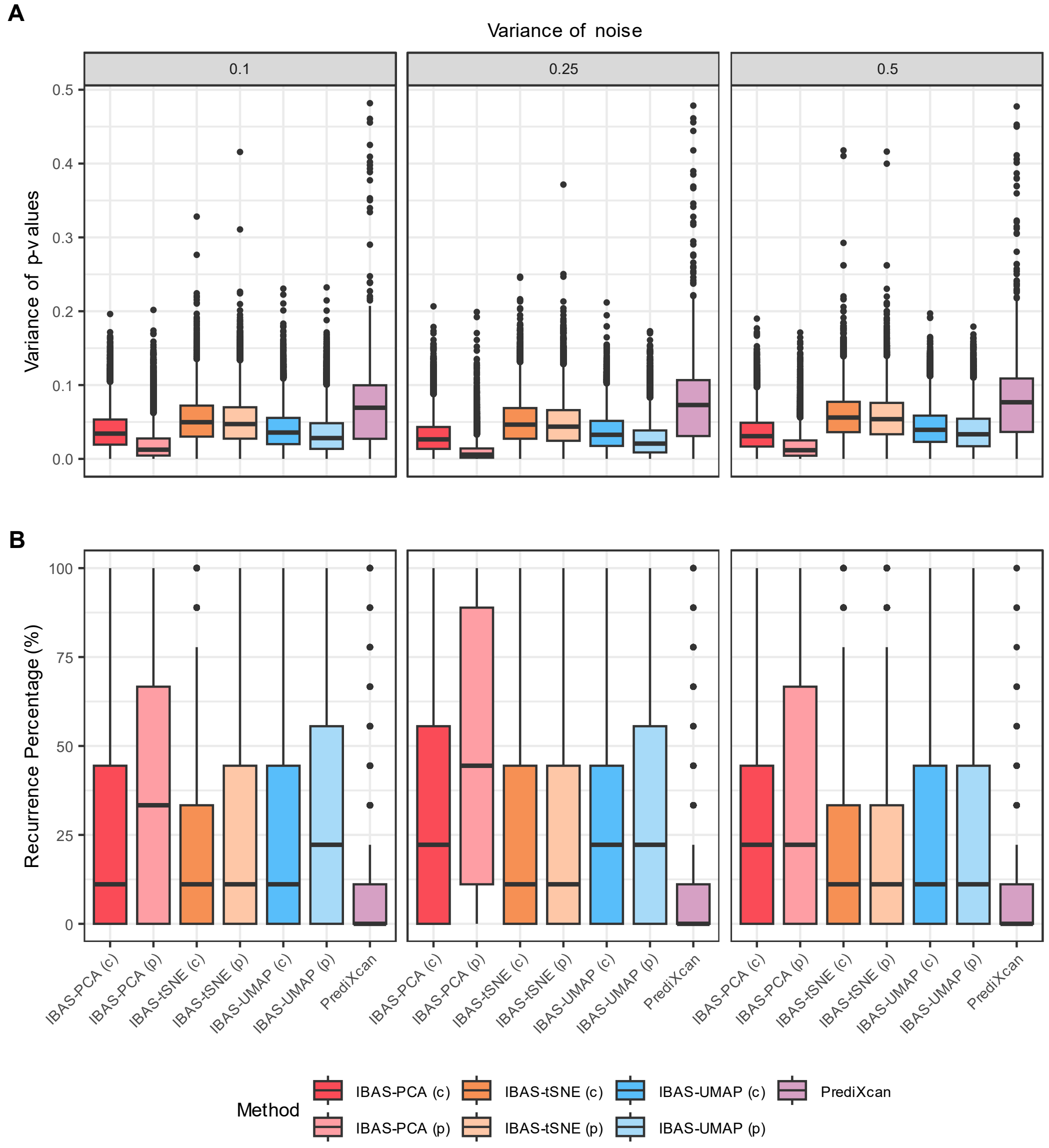

Simulations were conducted by perturbing gene expression values of each gene by percentage ranges (0.1, 0.25 and 0.5) in the reference data. 10 perturbed expression datasets (replicates) were generated for each of the noise levels and variability of results for case-control association with the Wellcome Trust Case Control Consortium Crohn's Disease cohort was evaluated using both IBAS and PrediXcan. PCA, t-SNE and UMAP dimensionality reduction methods as well as p-value based (p) and coefficient based (c) interaction association weights were tested in the case of IBAS. **A)** Indicates the variance of observed p-values in genes identified as significant ($p < 0.05$) in at least one replicate while **B)** indicates the recurrent significance of genes identified as significant in at least one replicate.

**Appendix Figure S4 – Simulated Perturbations reveal robustness of IBAS to noise in reference data (HT).**

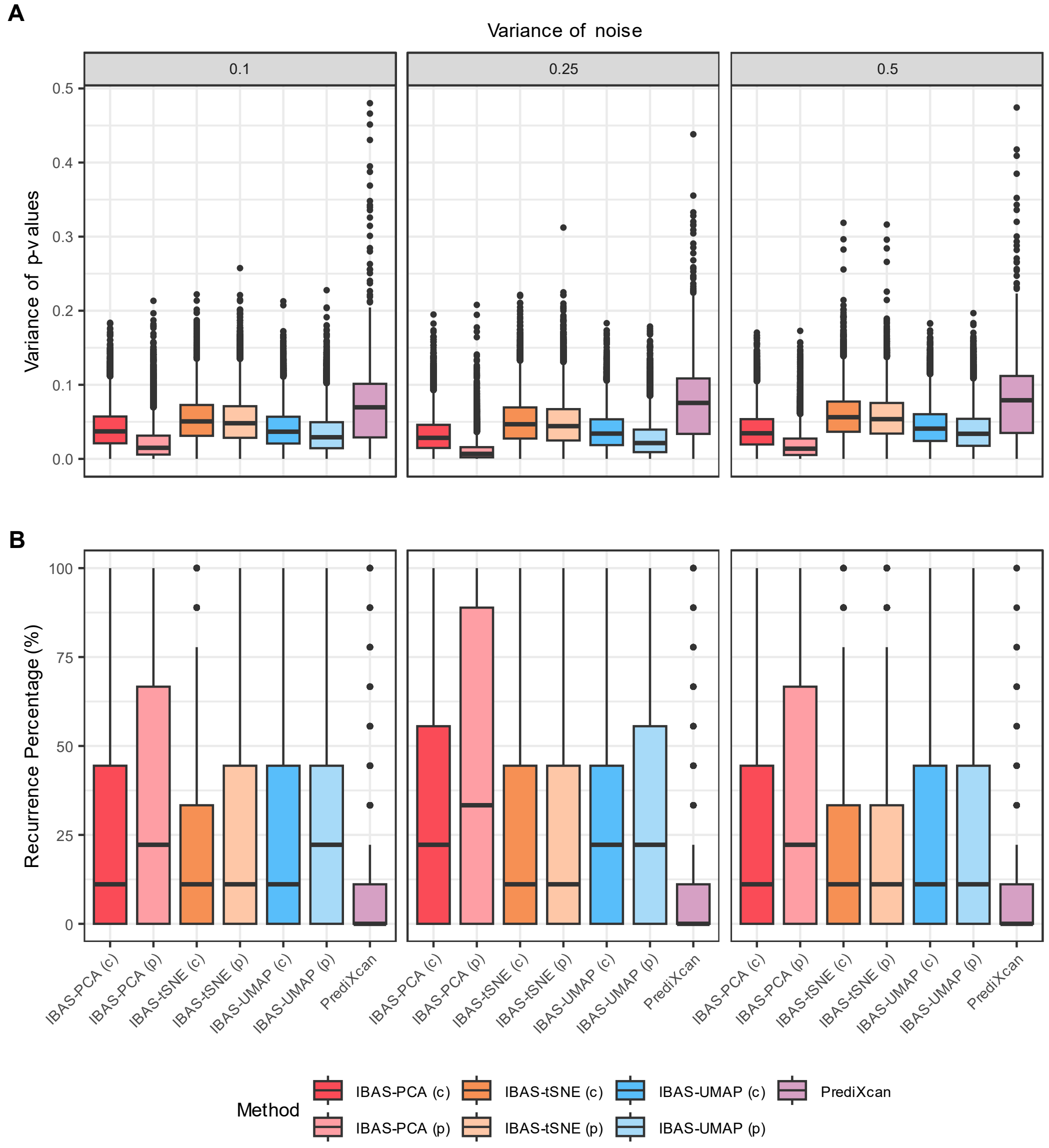

Simulations were conducted by perturbing gene expression values of each gene by percentage ranges (0.1, 0.25 and 0.5) in the reference data. 10 perturbed expression datasets (replicates) were generated for each of the noise levels and variability of results for case-control association with the Wellcome Trust Case Control Consortium Hypertension cohort was evaluated using both IBAS and PrediXcan. PCA, t-SNE and UMAP dimensionality reduction methods as well as p-value based (p) and coefficient based (c) interaction association weights were tested in the case of IBAS. **A)** Indicates the variance of observed p-values in genes identified as significant ($p < 0.05$) in at least one replicate while **B)** indicates the recurrent significance of genes identified as significant in at least one replicate.

**Appendix Figure S5 – Simulated Perturbations reveal robustness of IBAS to noise in reference data (RA).**

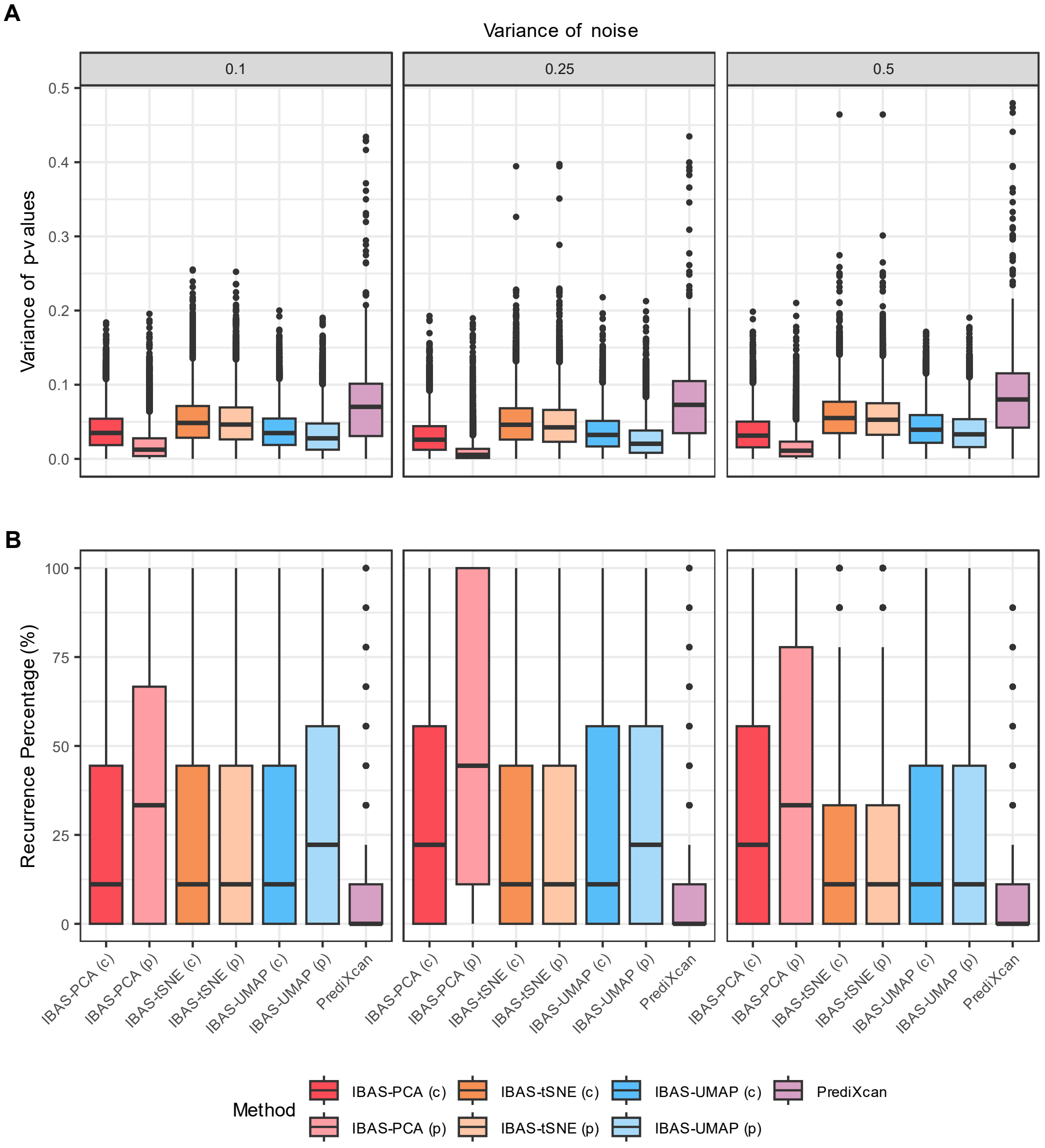

Simulations were conducted by perturbing gene expression values of each gene by percentage ranges (0.1, 0.25 and 0.5) in the reference data. 10 perturbed expression datasets (replicates) were generated for each of the noise levels and variability of results for case-control association with the Wellcome Trust Case Control Consortium Rheumatoid Arthritis cohort was evaluated using both IBAS and PrediXcan. PCA, t-SNE and UMAP dimensionality reduction methods as well as p-value based (p) and coefficient based (c) interaction association weights were tested in the case of IBAS. **A)** Indicates the variance of observed p-values in genes identified as significant ($p < 0.05$) in at least one replicate while **B)** indicates the recurrent significance of genes identified as significant in at least one replicate.

**Appendix Figure S6 – Simulated Perturbations reveal robustness of IBAS to noise in reference data (T2D).**

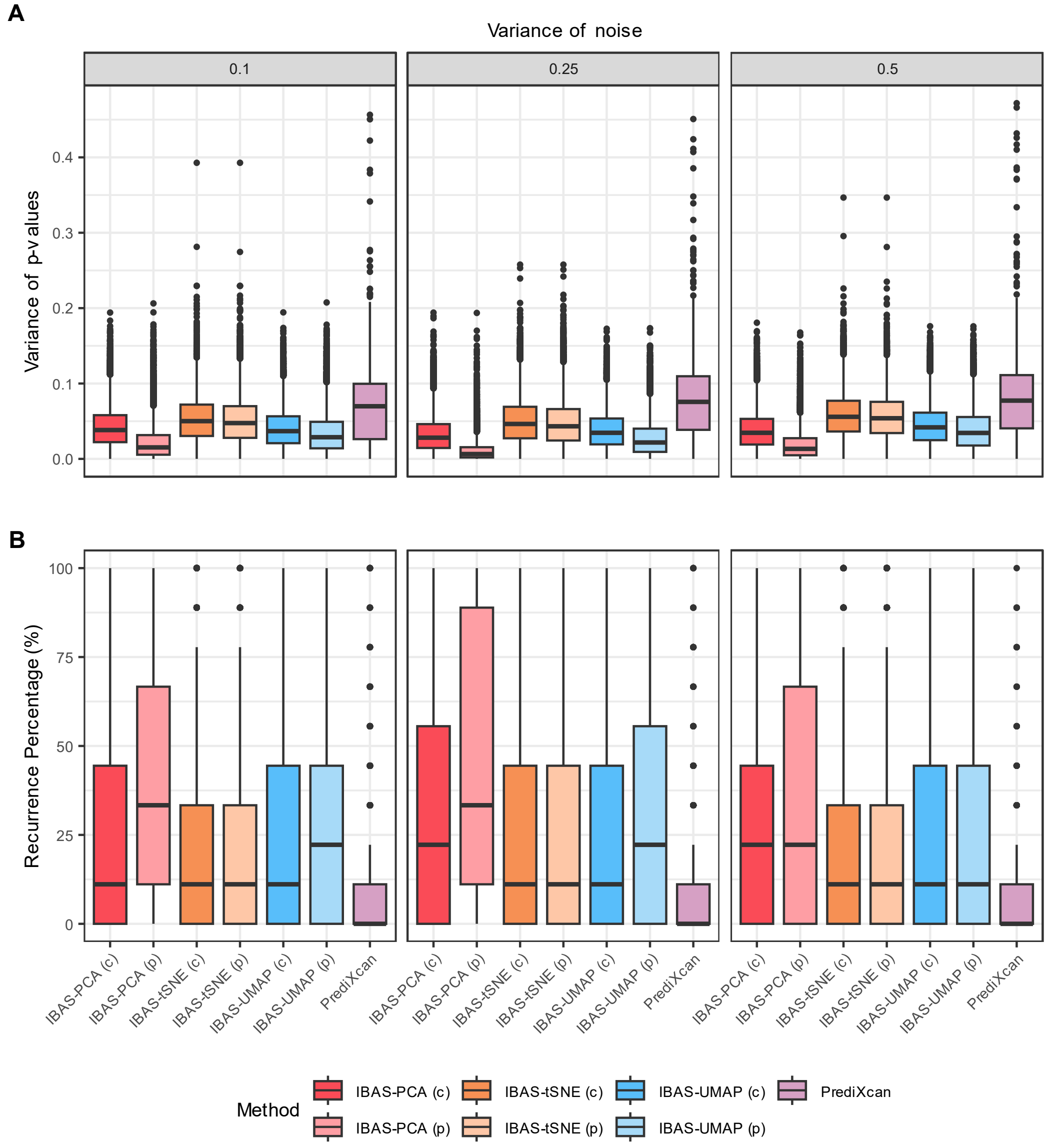

Simulations were conducted by perturbing gene expression values of each gene by percentage ranges (0.1, 0.25 and 0.5) in the reference data. 10 perturbed expression datasets (replicates) were generated for each of the noise levels and variability of results for case-control association with the Wellcome Trust Case Control Consortium Type 2 Diabetes cohort was evaluated using both IBAS and PrediXcan. PCA, t-SNE and UMAP dimensionality reduction methods as well as p-value based (p) and coefficient based (c) interaction association weights were tested in the case of IBAS. **A)** Indicates the variance of observed p-values in genes identified as significant ($p <$ 0.05) in at least one replicate while **B)** indicates the recurrent significance of genes identified as significant in at least one replicate.

**Appendix Figure S7 – Evaluation of genes associated with disease uncovered by IBAS-UMAP.**

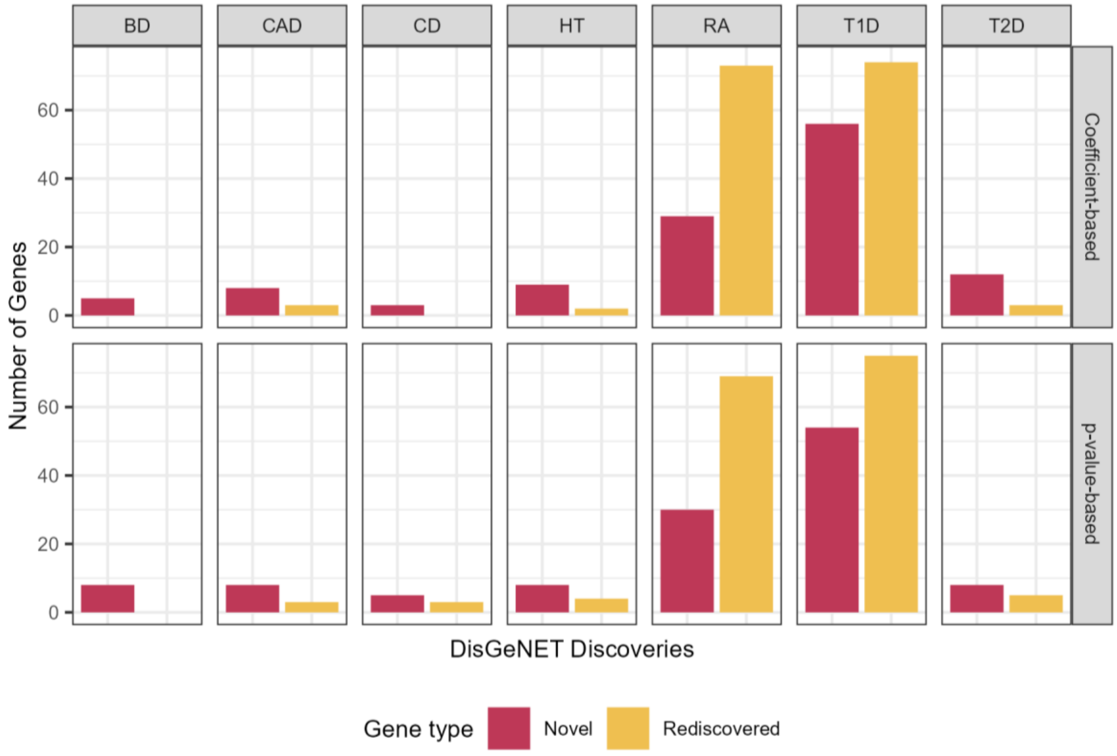

Genes identified as associated with each of the 7 diseases (columns) in the WTCCC dataset (with GTEx used as reference) across both coefficient-based and p-value based (rows) for the UMAP data-bridge were annotated as "Rediscovered" if found as previously associated with the same disease in the DisGeNET database. Remaining genes are annotated as "Novel".

**Appendix Figure S8 – Evaluation of genes associated with disease uncovered by IBAS-tSNE.**

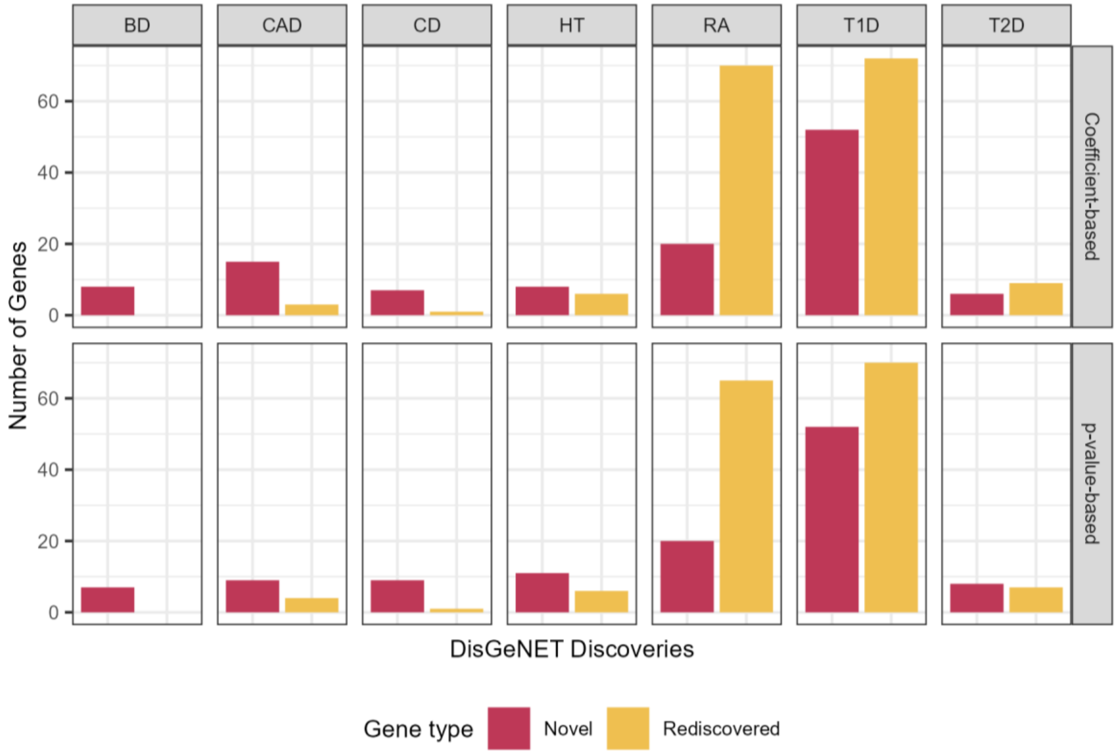

Genes identified as associated with each of the 7 diseases (columns) in the WTCCC dataset (with GTEx used as reference) across both coefficient-based and p-value based (rows) for the t-SNE data-bridge were annotated as “Rediscovered” if found as previously associated with the same disease in the DisGeNET database. Remaining genes are annotated as “Novel”.
